# Two-photon patterned photostimulation with low-power, high-efficiency and reliable single-cell optogenetic control

**DOI:** 10.1101/2022.01.08.475477

**Authors:** Yifan Wang, Yao Zheng, Yongxian Xu, Rongrong Li, Yameng Zheng, Jiajia Chen, Xiaoming Li, Hailan Hu, Shumin Duan, Wei Gong, Ke Si

**Author notes:** these authors contributed equally to this work (Yifan Wang, Yao Zheng).

## Abstract

Two-photon optogenetics enables selectively stimulating individual cells for manipulating neuronal ensembles. As the general photostimulation strategy, the patterned two-photon excitation has enabled millisecond-timescale activation for single or multiple neurons, but its activation efficiency is suffered from high laser power due to low beam-modulation efficiency. Here, we develop a high-efficiency beam-shaping method based on the Gerchberg-Saxton (GS) algorithm with spherical-distribution initial phase (GSSIP) to reduce the patterned two-photon excitation speckles and intensity. It can well control the phase of shaped beams to attain speckle-free accurate patterned illumination with an improvement of 44.21% in the modulation efficiency compared with that of the traditional GS algorithm. A combination of temporal focusing and the GSSIP algorithm (TF-GSSIP) achieves patterned focusing through 500-μm-thickness mouse brain slices, which is 2.5 times deeper than the penetration depth of TF-GS with the same signal-to-noise ratio (SNR). With our method, the laser power can be reduced to only 55.56% of that with traditional method (the temporal focusing with GS, TF-GS) to reliably evoke GCaMP6s response in C1V1-expressing cultured neurons with single-cell resolution. Besides, the photostimulation efficiency is remarkably increased by 80.19% at the same excitation density of 0.27 mW/μm^2^. This two-photon stimulation method with low-power, reliable and patterned illumination may pave the way for analyzing neural circuits and neural coding and decoding mechanism.

## Introduction

Optogenetics, using light to control neural activity with exogenously expressed light-sensitive proteins ^1, 2^, is a powerful neuroscience approach for dissecting the working mechanism of the brain ^3, 4^. In particular, temporally precise control of one or several individually selected cells is the key to manipulate the neural ensembles ^5-7^. Conventional optical realizations of driving neuronal activity have been achieved with visible light (single-photon excitation) ^8-10^, which fires the opsin-expressing neurons simultaneously within the field of view and cannot permit fast and precise optical manipulation of neuronal photostimulation at the single-cell level. It is still challenging to achieve cellular-resolution photoactivation for target neurons ^6, 11, 12^.

The holographic single-photon photostimulation approach offers efficient activation of cells with spatially selective illumination ^13, 14^. However, because of the light absorption and scattering in biological tissue^15, 16^, the realization of a near-cellular resolution relies on a micro-objective-coupled fiber bundle implanted in mouse brain ^17^, which induces acute^18^ and chronic^19^ tissue damage. Two-photon optogenetics, which largely mitigates the effect of scattering^20^ and provides single-cell-resolution photostimulation, has been achieved by the scanned method^21-23^ or the patterned method^24-26^. For extensively used opsins such as C1V1 and ChR2, the two-photon multi-foci scanning method increases the fraction of the activated cell membrane ^27^. However, such a strategy suffers from the loss of temporal resolution. Parallel-patterned light-targeting strategies have been proposed to stimulate opsins with millisecond and microscale resolution, which generate patterned illumination by sculpting light according to the morphology of neurons ^24, 28, 29^.

Two major patterned two-photon optogenetic systems, the combination of temporal focusing and generalized phase contrast (TF-GPC) ^24, 30^ or computer-generated holographic technology (TF-CGH) ^23, 29^, have been developed using spatial light modulators (SLM) or digital micromirror-based devices (DMD) to realize phase modulation. The TF-GPC system generates uniform-intensity patterned illumination from the interference of the binary-phase-modulation light with a synthesized reference wave (SRW), which is produced by a phase-contrast filter (PCF) ^31, 32^. TF-GPC has been used to fire action potentials in mouse acute cortical slices ^24, 30^, but the background noise of the TF-GPC illumination is so strong ^33, 34^that it limits the photoactivation efficiency. An alternative approach is TF-CGH based on the Gerchberg-Saxton (GS) algorithm (hereinafter called TF-GS for distinction). Compared with TF-GPC, TF-GS has a compatible optical configuration without the PCF and thus has been commonly used in two-photon optogenetics ^6, 29, 35^. However, the spatial intensity of illumination patterns based on the TF-GS system has conspicuous fluctuations (speckles) ^28, 36^, thus further limiting the improvement of photoactivation efficiency. The above consideration suggests that the conventional patterned two-photon excitation for simultaneous multi-cell photostimulation requires higher laser intensity ^2^ because the laser power needs to reach 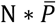 for stimulating N cells, where 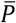 represents the average laser power for driving a single neuron. An intriguing question is whether it is possible to develop a high-efficiency beam-shaping method to decrease 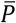 in the patterned two-photon photostimulation, which improves energy utilization, thus reducing the required laser power.

Here, we propose a method combining the Gerchberg-Saxton algorithm with spherical initial phase and temporal focusing two-photon microscopy (TF-GSSIP). TF-GSSIP optimizes the phase mask calculation method through prior conditions, which alters characteristics of random-mutation phase masks in typical CGH algorithms and fundamentally eliminates holographic speckles. It can precisely perform superior spatiotemporal patterned illumination with the modulation efficiency increasing by 44.21% compared with the GS algorithm. Besides, we present the scattering resilience properties of TF-GSSIP by scattering the patterned foci with 150-μm-thickness mouse skulls or 500-μm-thickness mouse brain slices. Compared to the TF-GS, the penetration depth of brain slices reaches a factor of 2.5 via the TF-GSSIP at the same SNR of patterned focusing. The result shows that TF-GSSIP enables to maintain the designed spatial excitation profiles and increase the excitation intensity to 201.61% of that in TF-GS using the same laser power. Finally, we set up a two-photon optogenetic system to confirm the high-efficiency photoactivation of TF-GSSIP in cultured cells. The TF-GSSIP system allows firing a significant increase of GCaMP6s fluorescence in C1V1-expressing neurons with single-cell resolution and ΔF/F increasing by 80.19% compared with TF-GS at the power density of 0.27 mW/μm^2^. For reliably evoking calcium response, the photostimulation power of TF-GSSIP is as low as 0.10 mW/μm^2^, only 55.56% of the laser energy used in TF-GS.

## Results

### GSSIP algorithm for High-diffraction-efficiency Beam-shaping

The GSSIP algorithm designs speckle-free patterned illumination tailored to the morphology of neurons to cover the target cell. In contrast to typical CGH algorithms that use random initial phase, the GSSIP algorithm to constrain the continuity of the phase mask (*φ*_*in*_). For scanless two-photon excitation of opsins, the general optical system for CGH converts incident light A_0_ on the input plane into a target intensity distribution A_*out*_ on the focal plane by phase modulation ^36^ (Extended Data Fig. 1a), so to a great extent, the photoactivation efficiency of TF-CGH is related to the phase mask. In the typical CGH system, the phase value of modulated beams between adjacent sampling points is close or more than π, which makes beams interfere to form holographic-speckle illumination on target neurons (Extended Data Fig. 1b) and affects holographic photoactivation effects. The GSSIP algorithm follows the general operation of the GS algorithm, which can be divided into two steps: initial values setting (A_in_ and *φ*_*in*_) and an iterative Fourier transform loop (IFTL), and finally outputs the predicted illumination pattern and the phase mask (Fig. 1a). The key of the GSSIP algorithm is the optimization of initial values, which are defined as a spherical-distribution initial phase related to the target illumination pattern and a Gaussian-distribution incident light field. According to the principle of the fast Fourier transform (FFT), the setting can maintain the phase continuity of adjacent pixels in the phase mask during IFTL. In the IFTL, after each iteration, the phase spectrums are retained, while the amplitude spectrums of the input light field (A_in_) and the output light field (A’) are replaced by A_0_ and A_out_, respectively. Combining the spherical initial phase with other quadratic phase distributions, such as linear-gradient phase term for off-axis beam shaping and conical phase term for hole beam shaping ^36^, will not affect the phase continuity of the phase mask. Therefore, the GSSIP algorithm can design arbitrary two-dimensional (2D) uniform (speckle-free) illumination patterns with rapid convergence (representative examples in Extended Data Fig. 1c and Extended Data Fig. 1d).

**Figure 1.**
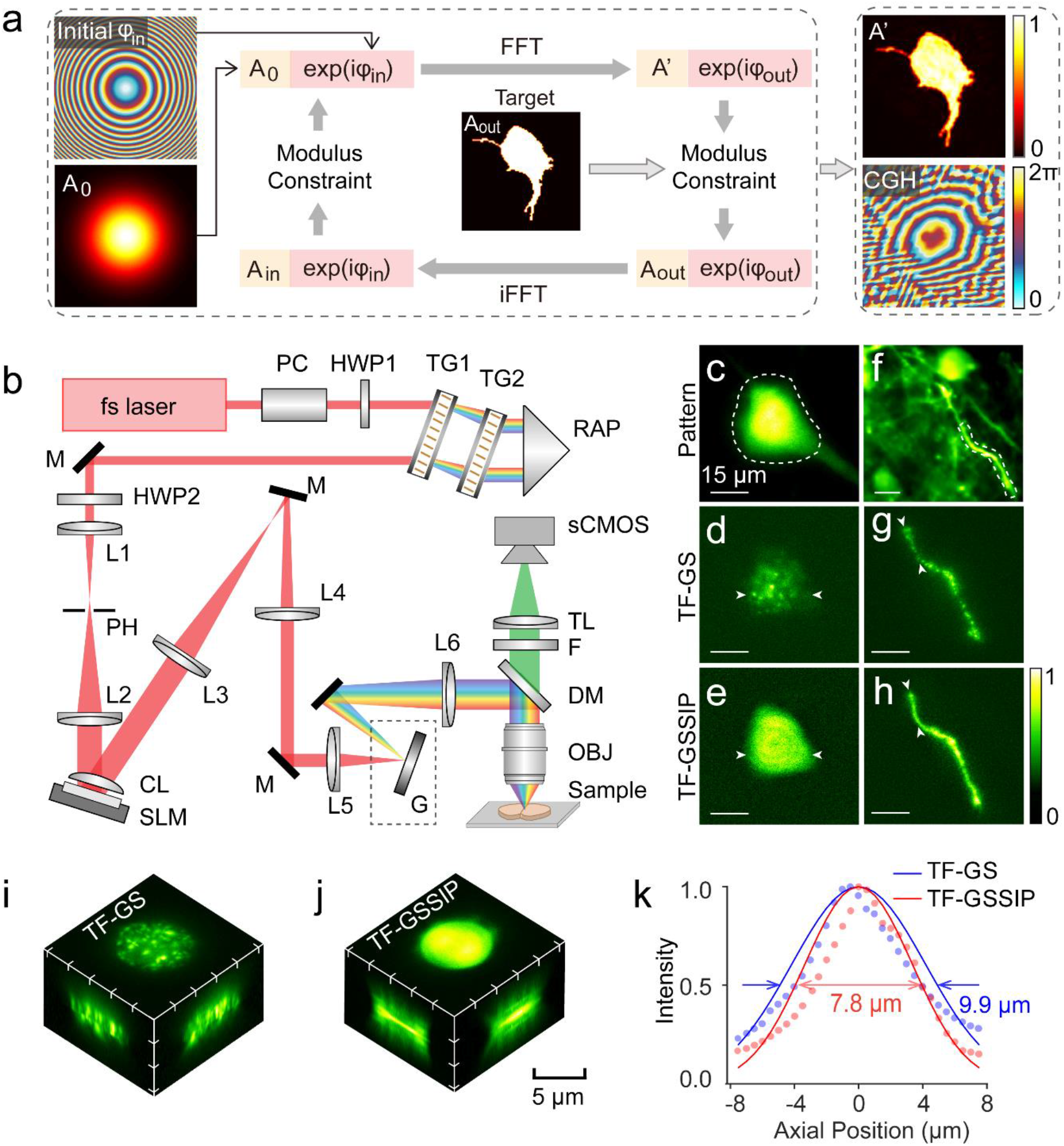
Diagrams of the algorithm and experimental setup of TF-GSSIP. **a** Flow diagram of the GSSIP algorithm. The differences between the GSSIP and GS algorithms are initial phase (*φ*_in_) and input beam (A_in_). **b** The optical set-up for TF-GSSIP. PC, Pockels cell; HWP, half-wave plate; TG, transmission grating; RAP, right-angle prism; M, mirror; L, lens; PH, pinhole; CL, cylindrical lens; SLM, spatial light modulator; G, blazed grating; DM, dichroic mirror; OBJ, objective; F, Filter; TL, tube lens. **c, f** Target patterns, cell body in Figure. c and a dendrite in Figure. f, based on a widefield fluorescence image of the brain slice (200-μm thickness, thy1-GFP mouse) and a two-photon scanning image of GCaMP6s-expressing neurons (awake mouse with an open-skull window). Scale bar, 15 μm. **d, e** Images of the neuronal soma by two-photon excitation of the target cell by TF- GS (Figure. d) and TF-GSSIP (Figure. e) two-photon excitation (brain slice, λ_exc_ = 920 nm; objective 20×, 1.0 NA). The intensity profiles along white arrows in **d, g, e, h** are shown in Extended Data Fig. 3a and Extended Data Fig. 3b) **g, h** Images of the dendrite by TF-GS (Figure. g) and TF-GSSIP (Figure. h) two-photon excitation of a thin fluorescent layer (Rhodamine B, λ_exc_ = 920 nm; objective 20×, 1.0 NA). **i, j** 3D projections of an 8-μm-diameter spot respectively generated by TF-GS (Figure. i) and TF-GSSIP (Figure. j). Scale bar, 5 μm. (Rhodamine B, λ_exc_ = 920 nm; objective 20×, 1.0 NA). **k** Normalized axial profile of the integrated fluorescence intensity of the 8-μm-diameter holographic spot by TF-GS (blue curve) and TF-GSSIP (red curve).

In contrast to other uniform CGH algorithms, the GSSIP algorithm shows a remarkable improvement in the performance of the designed illumination patterns. The region partition methods, such as mixed-region amplitude freedom (MRAF) algorithm ^36^ and double-constraint GS algorithm (DCGS) ^37^, are surrounded by extremely strong background noise which may perturb other neurons adjacent to the target neuron (Extended Data Fig. 2a-b). GSW-PC (weighted GS algorithm with phase- controll) algorithm developed for speckle-free 1D CGH ^34^ exhibits edge contour characteristics for 2D CGH (Extended Data Fig. 2c) and cannot achieve patterned illumination of the whole cell body. The phase-loop optimization algorithm optimizes the initial phase through multiple IFTAs and calculates the phase mask for uniform CGH (Extended Data Fig. 2d) ^38^. The diffraction efficiency and accuracy of the illumination pattern are similar to the result of the GSSIP algorithm (0.467 vs 0.468), but the multiple IFTAs extend the algorithm running time to dozens of seconds (11.60 s vs 0.88 s) (Extended Data Fig. 2j), which is detrimental to the switching of light-targeting modes in patterned two-photon optogenetics. Compared with the GS algorithm (Extended Data Fig. 2e), the GSSIP algorithm has made the illumination patterns 2 times more accurate (RMSE = 0.26 vs RMSE = 0.12) and 1.44 times higher modulation efficiency (0.32 vs 0.47) within the same running time. For star-shaped patterned illumination, the GSSIP algorithm achieves high-efficiency beam shaping (RMSE = 0.12, Modulation Efficiency = 0.47, Uniformity = 0.79) within an ms-level running time, which indicates this method can photostimulate target neurons with lower incident excitation density in two-photon optogenetics.

### TF-GSSIP system for High-efficiency Patterned 2P Excitation

To the imitate morphology of neurons, we utilize this algorithm combined with temporal focusing and characterize the system performance. The TF-GSSIP uses the same optical design in previously reported TF-CGH systems including temporal focusing and a phase modulation device (spatial light modulator, SLM) ^6, 28^, which constrains the axial resolution of excitation patterns (Fig. 1b). To generate and switch precise excitation patterns, any adjacent pairwise lenses placed between the SLM and the objective are conveniently fine-tuned to the telescope structure. A blazed grating (G) is placed at the front focal plane of lens L6 for temporal focusing ^39^. To simulate the fluorescence detection in the mammalian brain, we design the reflection fluorescence collection light path. In the optical configuration, the center of the field of view is the beam focus point when all pixels of the SLM are set to zero. It should be noted that the SLM under the working wavelength is calibrated in advance. According to the calibrated look-up-table (LUT), the phase mask generated by the GSSIP algorithm is loaded on the SLM while the TF-GSSIP system is working.

We demonstrate two examples of TF-GSSIP illumination visualized by exciting a brain slice (thy1-GFP) and a thin fluorescent layer (Rhodamine B): a neuronal soma and a dendrite. Excitation shapes are tailored to the geometry of neuronal fluorescence images (Fig. 1c, f). Using the same excitation intensity, TF-GSSIP achieves homogeneous-intensity and sharp-edged two-photon excitation (Fig. 1e, h, normalized fluorescence intensity profiles along the white arrows are shown in Extended Data Fig. 3a,b), which increases the signal-to-noise ratios (SNR) of the soma of 254.41% and the axon of 127.11% respectively compared with that of the TF-GS results (Fig. 1d, g). Moreover, the modulated wavefront of TF-GSSIP has a continuously spatial output phase mask (Extended Data Fig. 3c), which provides the optimal conditions for temporal focusing to constrain the axial resolution ^40^.

To characterize the effect of spatiotemporal focusing, the 3D focus fluorescence volumes of an 8-μm-diameter holographic spot are recorded on a thin fluorescent layer (Rhodamine B) by moving the axial motorized translation stage (Thorlabs, ZFM2020) (Fig. 1i, j). The axial range of the illumination pattern is defined as the full width at half maximum (FWHM) of the axial integrated intensity, which is calculated from the different planes of the optical stacks and shown in Fig. 1k. Due to the light energy is concentrated in the focus area, the TF-GSSIP system has a 7.8-μm axial resolution, while TF-GS has a value of 9.9 μm. The axial resolution of TF-GSSIP is smaller than the size of neurons (10-20 μm).

It is worth noted that the TF-GSSIP achieves the high two-photon excitation intensity and optimal axial constraint, which enables to generate powerful and uniform patterned illumination to efficiently stimulate target cells without crosstalk between adjacent neurons.

### A Comparison Between TF-GSSIP and Conventional Patterned 2P Optogenetic System through Biological Tissue

The main advantage of the temporal-focusing two-photon system is near-diffraction-limit focusing despite being scattered by biological tissue ^41, 42^, and patterned two-photon illumination also has this characteristic^28, 30^. We test the propagation of a 15-μm- diameter holographic spot through two kinds of scattering samples: 100∼500-µm-thickness mouse brain slices and 50∼150- μm-thickness mouse skulls, which are prepared from C57 mice (experimental configuration shown in Fig. 3a). Due to the low excitation efficiency of RhodamineB, the patterned focusing after propagating biological tissue is visualized by exciting a fluorescence slide with high cross-sections for two-photon excitation (Thorlabs, FSK2). And defocused fluorescence of the thick fluorescence slide cannot be excited premised on the high axis resolution of the system.

Under the same laser power, TF-GS used for conventional patterned two-photon excitation^6, 29^ fails to form a uniform holographic spot and demonstrates a rapid decrease in the intensity of the focused pattern as the thickness of brain slices is increases due to the strong scattering and absorption of the tissue (Fig. 2b, top row). And through a 500-μm brain slice, the envelope of the intensity profile is no longer visible (Extended Data Fig. 4a). In contrast, a recognizable focused holographic spot with the shape of the target pattern is maintained in brain slices up to 500 μm thickness (Fig. 2b, bottom) because TF- GSSIP shapes the illumination pattern with high modulation efficiency. Fig. 2c shows the SNR of the holographic imaging across a series of brain slice thicknesses. The TF-GSSIP system significantly enhances the SNR of holographic imaging across tissue thicknesses from 0 to 500 μm, as high as 201.61% at a slice thickness of 500 μm. And at the SNR of ∼2 dB, the TF-GSSIP increases the penetration depth to 500 μm, as a factor of 2.5 compared to that of the TF-GS. In addition, the shape of the holographic spot, generated by the TF-GSSIP system, appears strong robustness to scattering (Fig. 3d). Using the spatially continuous phase mask offers a good optical condition for the temporal focusing, so the TF-GSSIP system achieves less than 1 μm change in the diameter of the patterned focus. On the contrary, the diameter of holographic spots generated by TF-GS broadens from 15.38±0.018 μm at a slice thickness of 0 μm to 21.47±1.90 μm at a slice thickness of 500 μm. We next explore the effect of tissue scattering on the uniformity of the holographic illumination. Because the scattering specimen caused the beam to diverge, the uniformity of the holographic spots is increasing with the thicker brain slice. The uniformity of TF-GS holographic imaging even exceeds that of the TF-GSSIP in 500-μm-thickness brain slices owing to light absorption by the tissue. The light intensity has an exponential decrease through the scattering medium, so the gap between the high-intensity sites and the low-intensity sites of the speckle illumination pattern is reduced with the increase of the transmission depth. However, the uniformity of TF-GS comes at the cost of light energy, which is not desirable in optogenetic experiments. The light scattering absorption of biological tissues inevitably affects the uniformity of the patterned foci, but the ability of TF-GSSIP to maintain high excitation intensity and precision is significantly important for selectively exciting neurons.

**Figure 2.**
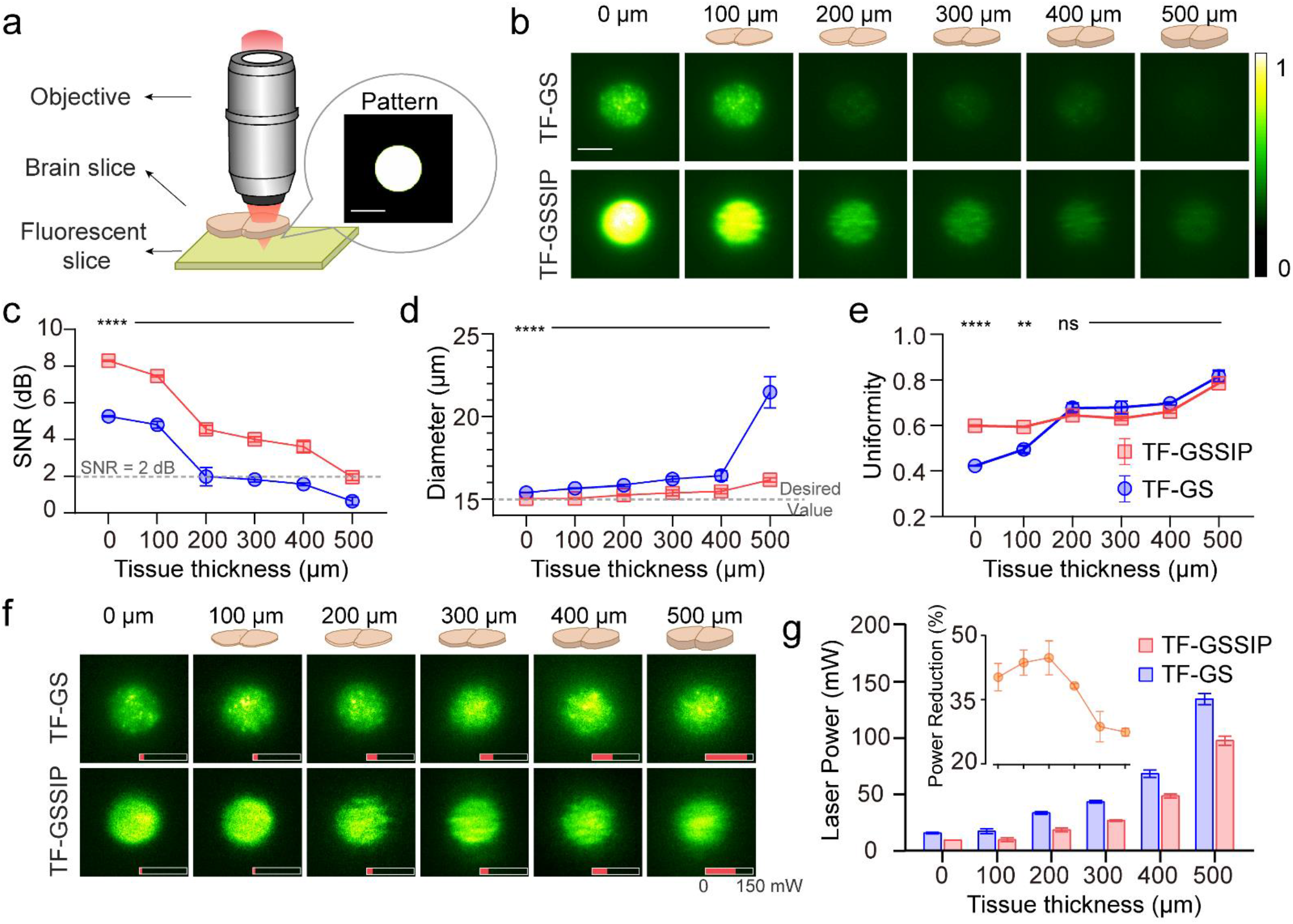
A comparison of TF-GSSIP patterned focusing and conventional patterned focusing. **a** The configuration for measuring the penetration depth in the scattering specimen. Sculpted beams focus on the fluorescent slide (FS) after propagating through brain slices. **b** Two-photon fluorescence images of a 15-μm-diameter holographic spot generated by TF-GS (top) and TF-GSSIP (bottom) after passing through fixed brain slices (0, 100, 200, 300, 400, and 500 μm) at the laser power of 0.10 mW/μm^2^. **c-e** SNR (Figure c), Diameter (Figure d), and Uniformity (Figure e) of the TF-GS and TF-GSSIP patterned foci as a function of tissue thickness. **f** Two-photon fluorescence images with the SNR of 1.76 dB at different laser power (shown by the red bar) generated by TF-GS (top) and TF-GSSIP (bottom) after passing through fixed brain slices with different thicknesses. **g** Statistical results of the laser power used for patterned two-photon excitation with the same intensity in TF-GS (blue) and TF-GSSIP (red). The inset shows compared with TF-GS, the reduced laser power is saved by TF-GSSIP in different brain slice thicknesses. Error bars in **c-e and g** represent the SD of five measurements taken at different locations. (λ_exc_ = 1064 nm; objective 40×, 0.8 NA)

**Figure 3.**
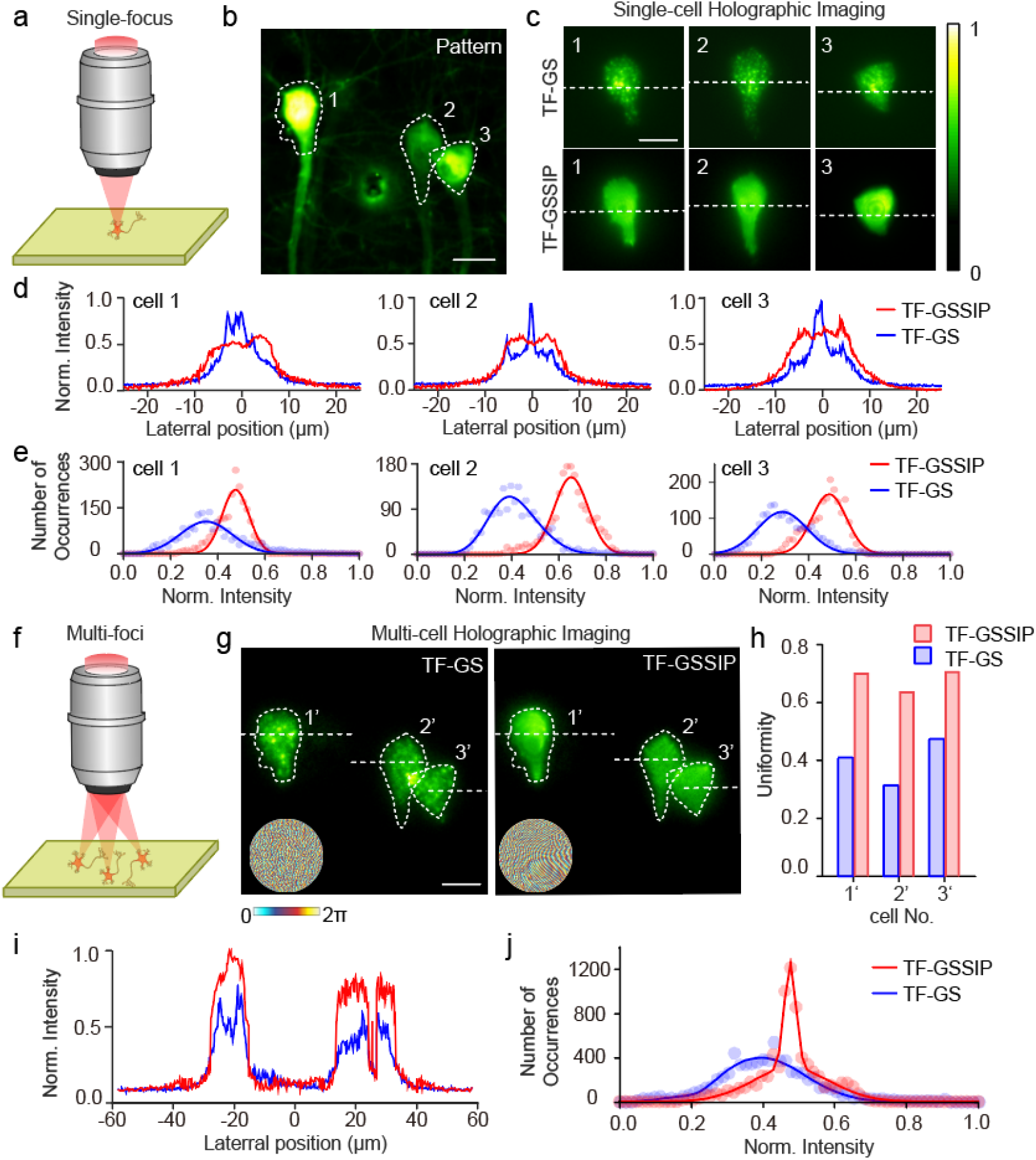
Single- and Multi-cell 2P excitation. **a** Schematic diagram of single-cell excitation. **b** A two-photon scanning imaging of a brain slice (thy1-GFP mouse), and cell bodies are selected as ROIs (white dotted line). **c** Three neuron-shaped fluorescence images by two-photon excitation of FSK2, which are generated by TF-GS (top) and TF-GSSIP (bottom). **d** Single-cell normalized fluorescent intensity profiles along the white dashed lines. **e** Histograms of ROI in the single-cell images. In **d** and **e**, the three results from left to right respectively present the features in neuron1, neuron2, and neuron3 (shown in Figure c). **f** Diagram of simultaneous multi-cell excitation. **g** Multi-cell fluorescent images generated by TF-GS (left) and TF-GSSIP (right), each cell lying in the same position as Figure b. Insets are phase masks designed by the GS and GSSIP algorithms. **h** The uniformity of three neuron-shaped illumination patterns designed by TF-GSSIP (red) and TF-GS (blue). **i, j** Normalized fluorescent intensity profiles along the white dashed lines and histogram of ROI in multi-cell fluorescent images. Scale bars, 15 μm. (λ_exc_ = 1064 nm; objective 40×, 0.8 NA)

The scattering experiment is also performed in the skull (C57 mice), which exhibits stronger scattering and absorption than brain slices. TF-GSSIP penetrates a 150-μm-thickness skull to form visible patterned two-photon excitation, while TF-GS fails to distinguish intensity profile at a 100-μm-thickness skull (Extended Data Fig. 4b). Therefore, the characteristics of spatially continuous phase mask and high modulation efficiency enable the TF-GSSIP to create accurately patterned two- photon excitation deep into a scattering specimen.

We next compare the laser power while TF-GSSIP and conventional patterned two-photon excitation generated the same intensity of holographic focusing through scattering samples. The laser power is recorded when the SNR of a 15-μm-diameter holographic spot is 1.76 dB (the ratio of the illuminated over the non-illuminated area to 1.5). TF-GS realizes visual two-photon patterned excitation by increasing the laser power, but notable spatial intensity fluctuations are invariably accompanied as the brain slice thickness is increasing (Fig. 2f, top). On the contrary, TF-GSSIP maintains speckle-free focusing patterns no matter how thick the brain slice is (Fig. 2f, bottom). Fig. 2g shows the difference in laser power between TF-GS and TF-GSSIP when the fluorescence intensities are the same. TF-GSSIP saves about 40% of laser energy in the 300-μm or thinner slices. Even in 400- or 500-μm brain slices, TF-GSSIP uses only 72.57% of the conventional patterned 2P excitation energy. Overall, the TF-GSSIP not only mitigates the influence of scattering but also saves 27.43%-44.76% of the laser energy corresponding to 0-500 μm brain slices, which indicates TF-GSSIP offers a precise and efficient patterned two-photon excitation in biological tissues.

### Accurate Patterned 2P Excitation for Single- or Multi-cell

The intensity of the illumination pattern generated by the 2P holography technology directly affects the efficiency of two-photon excitation, such as the optimal excitation is obtained when the sculpted beams just cover targeted neurons ^24^. TF-GSSIP can generate speckle-free patterned illumination with high diffraction efficiency to optimize the patterned two-photon excitation. In this section, we demonstrate TF-GSSIP generates single- or multi-target two-photon excitation, and compare them with the TF-GS.

The single-target patterned 2P excitation is presented to confirm that the high homogeneity and accuracy of TF-GSSIP are pattern-independent (optical diagram shown in Fig. 3a). Different excitation patterns are customized according to the geometry of cell bodies in a two-photon scanning fluorescence imaging (from a thy1-GFP mouse, Fig. 3b). Compared to the TF-GS (Fig. 3c, top), TF-GSSIP generates every neuron-shaped 2P excitation pattern with uniform fluorescence intensity and obvious profiles (Fig. 3c, bottom). Because of the absence of random speckle-noise, TF-GSSIP provides holographic imaging with as twice accuracy as TF-GS does (see Extended Data Table 1). In Fig. 3d, the normalized intensity profiles of three cells show that TF-GSSIP eliminates notable spatial intensity fluctuations in holographic 2P excitation. In addition, the SNRs of three neuron-shaped images generated by TF-GSSIP respectively increase to 131.56%, 150.99%, and 155.72% of that of TF-GS due to high diffraction efficiency. For uniformity, we calculate the normalized grayscale histogram of regions of interest (ROI) (Fig. 3e). The intensity fluctuations of more than 90% pixels in each neuron-shaped fluorescence image generated by TF-GSSIP are 0.24, 0.18, and 0.23, corresponding to TF-GS are 0.35, 0.36, and 0.35, respectively.

TF-GSSIP also maintains homogeneously high intensity independent of the number of targets. We deliver multi-cell 2P excitation results according to the generalized adaptive-additive (GAA) method ^43, 44^ (Fig. 3h), and the phase masks designed by the GS algorithm and the GSSIP algorithm are shown in Fig. 3h illustration. In agreement with the results of single-cell 2P excitation, the multi-cell 2P fluorescence images obtained by the TF-GSSIP have flat intensity profile (Fig. 3g, i) and higher accuracy (see Extended Data Table 1). The uniformity of three ROIs in the multi-target excitation produced by TF-GSSIP is 0.64, 0.70, and 0.71, respectively, which is 68.83%, 99.40%, and 47.67% higher than that of TF-GS (Figure 3i). Through the normalized gray-scale histogram fitting results, we find that TF-GSSIP offers multi-target 2P excitation with fluorescence intensity varying less than 0.08 in more than half of the pixels in the ROI, corresponding to more than 0.12 variation generated by TF-GS (Fig. 3g).

These results confirm that TF-GSSIP light patterning automatically creates speckle-free excitation shapes with high efficiency and high accuracy for single- or multi-target excitation. This is similar to the function of 2D patterned two-photon optogenetic devices, such as TF-GPC ^24^ or TF-GS ^29^, but TF-GSSIP has higher beam shaping efficiency to form stronger two- photon excitation.

### All-optical single-cell resolution stimulation and recording in Cultured Neurons

Patterned photostimulation is a spatiotemporal-flexibility light-targeting technology to reliably fire action potentials of neurons ^2, 45^. However, parallel- patterned illumination requires higher laser power than scan-based strategies ^2^. The TF-GSSIP, as a high-diffraction-efficiency beam shaping method, can significantly reduce the laser energy required to activate a single neuron. Thus, compared to TF- GS, TF-GSSIP realizes reliable photoactivation at low power density. To illustrate the performance of TF-GSSIP for cell photoactivation, we compared the photostimulation efficiency by measuring the peak value of calcium response during optical recordings and perturbations of neuronal activity. First, TF-GSSIP generates patterned illumination tailored to the morphology of target cells, which is used to stimulate dissociated rat cultured hippocampal neurons infected with adeno-associated virus (AAV) encoding GCaMP6s and C1V1-mCherry (Extended Data Fig. 5b). For functional imaging, we integrated an epi-fluorescence microscope in the TF-GSSIP system (shown in Extended Data Fig. 5a). In the integrated system, the wavelength of the laser source is switched to 1,064 nm for exciting C1V1.

To validate the single-cell-resolution photostimulation by TF-GSSIP, we selectively stimulate a single cell with 3s pulses at a rate of 0.043 Hz for 90 s (excitation intensity 0.10 mW/μm^2^, corresponding to 3.63 mW/cell), while recording the neuronal activity at a frame rate of 20 Hz with an illumination power of 0.12 mW/mm^2^ (Fig. 4a). With this protocol, TF-GSSIP and TF- GS lead to an increase in GCaMP6s fluorescence signal (Fig. 4a, top trace). TF-GSSIP not only evokes a more prominent calcium response compared with the TS-GS but also shows a significant difference between the target neuron and neighboring neurons (Fig. 4a, right). Thus, the TF-GSSIP has better specificity compared with the TF-GS. Moreover, the TF-GSSIP offers a higher photoactivation probability compared with the TF-GS under light power of 0.10 mW/μm^2^ (n = 9 cells, t test, *p = 0.032) (Fig. 4b). Reliable photoactivation under low laser power indicates that the uniform illumination pattern generated by TF- GSSIP covers more C1V1-expressing sites on the cellular membrane, and C1V1-mediated cation influx at each opsin-expressing site, eliciting a rapid increase of GCaMP6s fluorescence ^14^. Consequently, TF-GSSIP is a more reliable patterned two-photon photostimulation method than TF-GS.

**Figure 4.**
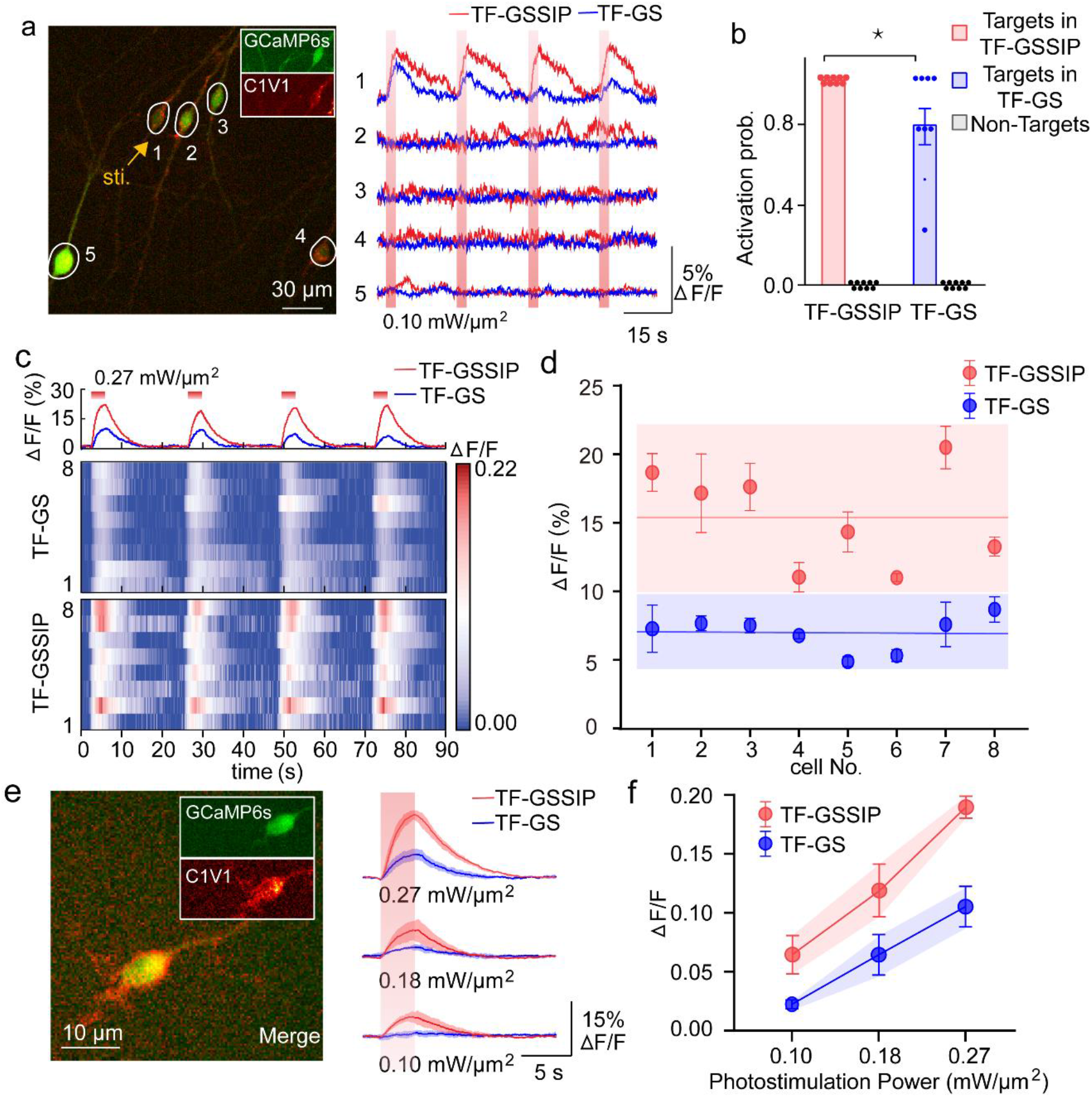
Comparison of photoactivation efficiency between conventional patterned photostimulation and TF-GSSIP. **a** Single- cell-targeted photostimulation by TF-GSSIP (red trace) and TF-GS (blue trace) system. A single neuron is targeted for C1V1 photostimulation at 1,064 nm with 3-s pulses (vertical red shaded bars) at a power of 0.10 mW/μm^2^ while GCaMP6s fluorescence is imaged at 488 nm (0.12 mW/mm^2^). Left, a maximum intensity projection wide-field fluorescence image of neurons, with a target neuron (yellow circle) and surrounding non-target neuron (white circle). Right, 2P Holographic- photostimulated responses from the corresponding target neuron (cell No.1) and nontarget neurons (cell No.2-5). Vertical red bars represents the light stimulation period. **b** Activation probability of targets and non-targets (black spots) stimulated by TF- GSSIP or TF-GS. Each red (TF-GSSIP) or blue (TF-GS) spot demonstrates the activation probability of the target cells in single- neuron photostimulation experiments (shown in **a**; 9 FOVs). All bar graphs depict mean ± S.D (p = 0.032, t test). **c** Heatmaps showing the target cell (n = 8) responses to holographic photostimulation by TF-GS (middle) and TF-GSSIP (bottom) system at the power of 0.27 mW/μm^2^. GCaMP6s responses to TF-GSSIP are higher than it to TF-GS. **d** The average evoked target-cell responses (mean ΔF/F ± S.D.; 4 trials for each cell; 0.27 mW/μm^2^ stimulation power; 3 s stimulation duration; 0.10-0.35 mW/mm^2^ imaging power). The minimum GCaMP6s responses to TF-GSSIP is higher than the maximum response of calcium signals to TF-GS. **e** Single-cell photostimulation by TF-GSSIP (red trace) and TF-GS (blue trace) system upon different stimulation power. Left, a merged wide-field fluorescence image of neurons coexpressing the opsin C1V1-mCherry (red) and calcium indicator GCaMP6s (green). Right, average calcium response to photostimulation at a power of 0.27 mW/μm^2^ (top), 0.18 mW/μm^2^ (middle) and 0.10 mW/μm^2^ (bottom) (mean ΔF/F ± SD, 4 trials; 3 s stimulation duration; 0.14 mW/ mm^2^ imaging power). **f** Average evoked calcium signal ΔF/F as a function of photostimulation power (detail data is shown in Figure S3).

Cultured neurons at 3-4 days after infection are selected as targets to explore the efficiency of patterned two-photon stimulation. Photostimulation of the target cell evokes GCaMP6s fluorescence response with the peak of fluorescence intensity dependent both on stimulation duration and power (Fig. 4c-f and Extended Data Fig. 5c-d). For a fixed stimulation power of 0.27 mW/μm^2^ with a duration of 3 s, the response of calcium imaging has a very short latency by TF-GSSIP photostimulation, and the intensity of the calcium response is significantly stronger than that of TF-GS (n = 8 cells, Fig. 4c). We measure the peak of calcium response of the target cells and find that TF-GSSIP increases GCaMP6s responses from 0.07 ± 0.02 to 0.15 ± 0.04 compared with the TF-GS photostimulation (Fig. 4d, n = 8 cells, t test, ****p < 0.0001). Besides, different laser powers are applied to stimulate target cells via TF-GSSIP and TF-GS (Fig. 4e). For a constant stimulation duration of 3 s, GCaMP6s responses increase linearly with photostimulation power both for TF-GS and TF-GSSIP (Fig. 4f). And a power density increases from 0.10 to 0.27 mW/μm^2^ lead to an increasing peak of GCaMP6s responses from 6.45±3.25% to 18.98±1.87% by TF-GSSIP, while if the same target neuron is stimulated by TF-GS, the peak only changes from 2.23±0.76% to 10.52±3.44% (n = 5 cells) (Fig. 4e-f, other neurons in Extended Data Fig. 5c). The results are similar under different stimulation durations. For 0.27 mW/μm^2^ stimulus power, the peak of GCaMP6s responses stimulated by TF-GSSIP increases monotonically from 2.64±0.62% for 1 s stimulation duration to 6.44±0.68% for 3 s stimulation duration, while in the TF-GS, the peak only increases from 1.88±0.54% to 3.47±0.94% with the stimulation duration ranging from 1 to 3 s (Extended Data Fig. 5d). The changing trends in calcium response with stimulation duration and power are consistent with the published paper about the number of action potential spikes triggered by TF-GS photostimulation ^29^. Of note, the TF-GSSIP 2P photostimulation also provides the target cell response twice as high as TF-GS upon different stimulus energy (Fig. 4f).

Altogether, these results indicate the TF-GSSIP system can realize light-targeting photostimulation with single-cell precision, and verify that the calcium responses to TF-GSSIP photostimulation are more obvious than TF-GS^6, 21, 28^, and TF- GPC^24, 30^. The main reason is that the holographic-photostimulation pattern of the TF-GSSIP system has a uniform distribution. Because of the scattered expression of light-sensitive proteins on the cellular membrane, the speckle-type holographic light spot of TF-GS fails to accurately cover all expressed opsin and the energy distribution is uneven. However, the uniform holographic illumination covers the entire cell indiscriminately, and all light-gated proteins at the focal plane take effect simultaneously during photostimulation. This high-sensitivity two-photon holographic photostimulation method enables to reduce patterned stimulation laser power for reliable activation of neurons from 0.18 mW/μm^2^ to 0.10 mW/μm^2^ (Fig. 4e, left and Fig. 4f).

## Discussion

Similar to the parallel-patterned light-targeting method that shows a great improvement over the serial-scanned two-photon microscope, TF-GSSIP provides superior-spatiotemporal-resolution patterned two-photon excitation. Moreover, TF- GSSIP sculpts beams with high modulation efficiency and great uniformity (speckle-free), which effectively reduces the laser power for firing calcium response. These advantages come without compromising its characteristic dynamically patterned excitation. TF-GSSIP achieves a shorter running time than TF-GS does, because the GSSIP algorithm converges rapidly by the initial value optimization (Extended Data Fig. 1e).

The TF-GSSIP allows precisely patterned focusing after propagating through the mouse skull and brain slices, proving the potential capability of photostimulation in vivo. This is due to the continuous distribution of spatial wavefront calculated by the GSSIP algorithm, improving the axial focusing performance of the spatiotemporal focusing system. Thus TF-GSSIP partly overcomes the scattering of biological tissue and maintains the excitation shape at a large depth. With the increase of tissue depth, the uniformity of the shape of the focal stimulation patterns inevitably deteriorates, and the intensity of the stimulus light plays a major role in the formation of effective photostimulation. The TF-GSSIP improves the efficiency of light modulation, so that the excitation light can still retain sufficient intensity at a deeper position, which improves the reliability of photostimulation and helps to realize the selective activation of single or multiple neurons in the deep brain.

Applying the TF-GSSIP combined with the GAA method to form 2D multi-target parallel two-photon excitation achieves enhanced SNR and high precision. We achieve accurately 3-neurons-shaped two-photon excitation, and more illumination patterns can be generated at the cost of reduced precision and uniformity because the crosstalk of numerous phase masks would smear the multi-target illumination. This tradeoff can potentially be circumvented by multiple mini-beams modulation, which means that the SLM is segmented into subregions ^46-48^, and each subregion has an individual phase mask designed for each target. Multi-targets addressing arbitrary 3D locations can also be realized if a second SLM is used in the TF-GSSIP for dedicated axial modulation, such as MTF-MS ^33^ or 3D-SHOT ^49^.

In comparison to current patterned two-photon optogenetic techniques, the TF-GSSIP excels in improving the photostimulation efficiency with a notably reduced laser power for photostimulation. Under a power density 44.44% lower than that required in conventional TF-CGH, TF-GSSIP enables to reliably fire the calcium response of targeted neurons with prominent fluorescent changes. Furthermore, this system can be combined with the two-photon scanning microscope expediently to explore the function of neuronal ensembles in vivo, which underlies a range of exploring the fine-scale organization of the brain ^5^.

## Acknowledgements

This work was supported in part by the National Natural Science Foundation of China (61735016, 81771877), Natural Science Foundation of Zhejiang Province (LR20F050002), Key R&D Program of Zhejiang Province (2020C03009, 2021C03001), Zhejiang leading innovation and entrepreneurship team (202099144), CAMS Innovation Fund for Medical Sciences (2019-I2M-5-057), Fundamental Research Funds for the Central Universities.

## Author contributions

K. Si and W. Gong conceived the concept and supervised the research. Y. Wang developed the GSSIP algorithm. Y. Wang and Yao Zheng performed all the experiments including designing and building the optical system. R. Li made the scattering samples and assisted with the scattering experiments. J. Chen assisted with the design of the temporal-focusing system. Y. Xu cultured in vitro hippocampus neurons and assisted with the photostimulation experiments. S. Duan, H. Hu, and X. Li guided the biological experiments and provided the mice. Y. Wang and Yao Zheng analyzed the data. All the authors discussed the results and wrote the manuscript.

## Methods

### Two-photon holographic microscopy for TF-GSSIP

The optical setup for two-photon holographic imaging or photostimulation is based on a customed multiphoton microscope system (Fig. 1a). The system realized the function switching of the imaging and photo-stimulation by adjusting the output wavelength of the tunable optical parametric oscillator (Chameleon Ultra II, COHERENT).

For fluorescence imaging, the green fluorescence of RhodamineB and GFP are excited at 920 nm with an 80 MHz repetition rate and 140fs pulse. The laser firstly passes through a pockels cell (Model 350-80-LA, Conoptics) for power adjustment and a linear film polarizer (P, Thorlabs, LPVIS050) for linear polarization. Then the dispersion compensation module, composed of two transmission gratings (TG1 and TG2, Wasatch Photonics, 600l/mm @ 900nm) and a right-angle prism (RAP, Union Optic, 3008010355), pre-compensates the beam dispersion from other optical dispersion elements in the system. After dispersion compensation, the beam is spatially filtered by two lenses (L1, Thorlabs, C560TME-B; L2, Golden Way Scientific, GL31-025-50-NIR) and a pinhole (P, Thorlabs, P30D) to fill the SLM active area with Gaussian distribution. And the polarization is optimized with a half-wave plate (HWP, Union Optic, WPA2420-650-1100) to achieve the maximum efficiency of the reflective SLM (Model P512-0785, Meadowlark Optics, 7.68×7.68 mm2 active area, 512×512 pixels). Two plano-convex cylindrical lenses (CL, Union Optic, CYX0015) are placed next to the SLM to eliminate zero-order diffracted light ^1^. After modulation by the holographic phase mask on SLM, the holographic beam is relayed by a telescope (L3, Golden Way Scientific, GL31-025-150-NIR; L4, Golden Way Scientific, GL31-025-100-NIR), and then focused on a reflection grating (G, Thorlabs, GR13- 0610) for temporal focusing by lens L5 (Golden Way Scientific, GL31-025-100-NIR). The first diffraction order is collimated by lens L6 (Golden Way Scientific, GL31-025-200-NIR) and reflected by a low-dispersion shortpass dichroic mirror (DM, Edmund, #86106), while the zero diffraction order is blocked by a beam block (B, Thorlabs, LB1). The holographic beam finally is focused as the designed illumination pattern on the sample by a 20 × /1.0 NA water immersion objective lens (OBJ, Olympus, XLUMPLFLN20XW). Emitted photons are collected by the same objective lens, and penetrate the filter (F, Semrock, FF01- 520/60-25). The two-photon fluorescence is detected by an sCMOS (Dhyana 400BSI V2, TUCSON).

2P photostimulation is performed at 1064nm with an 80 MHz repetition rate and 110fs pulse. Except for the output wavelength of the tunable optical parametric oscillator, the optical configuration for holographic photostimulation is consistent with the above-mentioned holographic imaging system. The holographic beam used for photoactivation is reflected by the dichroic mirror (DM1, Edmund, #86106), and focused by a 40 × /0.8 NA water immersion objective lens (OBJ, Nikon, CFI Apo NIR 40X W) to stimulate the target neuron. To record the calcium response of neurons, the fluorescence of GCaMP6s is observed by an epifluorescence-inverted microscope (Extended Data Fig. 5a). The warm white light from the halogen lamp (Olympus, IX3-RFA) only penetrates the blue light at about 488nm when passing through the filter (F1, Semrock, FF01-482/35). Then the beam is reflected by a long-pass dichroic mirror (DM2, Semrock, FF506-Di03) into the Nikon objective lens. The wide- field excited GCaMP6s fluorescence is collected by the same objective and finally detected by the sCMOS.

Phase masks generated by GSSIP or GS algorithm are loaded on the SLM using the manufacturer’s software development kit (SDK) and custom experimental control software with a calibrated look-up table (LUT) written in MATLAB. The registration of the photostimulation FOV and the wild-field imaging FOV is performed before each experimental session to ensure effective cell-targeted stimulation. The photostimulation system irradiates a thin fluorescent film (RhodamineB) with a calibration pattern. The film is imaged by the wide-field system, and a calibration pattern is used for the registration of the two systems.

### SLM calibration

There are two kinds of static errors related to the electric-addressed SLM performance, respectively called the global grayscale-phase mismatch and spatial nonuniformity ^2, 3^. The former is due to the inappropriate preset look-up table (LUT) for the SLM. The latter comes from the nonuniform electric drive ^4^ or the curvature of the backplane of the SLM ^5^. Moreover, the precision of the applied electric field transferred from the uploaded grayscale for the pixels also introduces errors. And the applied electric field is changing with illuminating wavelength. LUT is used to map grayscale values, linear to the applied electric field, to 0 to 2π phase shift. Therefore, calibration for an accurate LUT is required for improving the performance of the SLM ^6^.

The interference calibration procedure ^6^ is used to calibrate the LUTat 920 nm and 1064 nm required for the experiment. The collimated laser beam expands to fill the incident surface of the SLM. The SLM and the camera are located on the front and back focal planes of the lens, which are respectively used to modulate the phase and capture the interference fringes. The method takes effect based on a mask with two pinholes placed in the front of the reflective SLM. To obtain the best contrast in the fringe pattern, we customize a mask with two pinholes with a distance of 4.2 mm and a diameter of 0.5 mm. The SLM is divided into two regions, and each is multiplied by a pinhole, where the intensity of the incident light is the same. The modulation phase can be measured by remaining the grayscale of one half of the SLM at a constant grayscale, while gradually changing the grayscale of the other half of the SLM from 0 to 65535 by linear LUT (16bit modulation for Model P512- 0785). The relatively shifted phase, from the grayscale changes, can be calculated by the cosine formula, which is corresponded to the known grayscale one-to-one to establish the gray-phase relationship.

We next test whether the calibrated LUT is practical in our system. The process of the verification experiment is the same as the above calibration steps. The difference is that the phase corresponds to the grayscale on both sides of the SLM according to the calibrated LUT. If the phase offset changes linearly with the grayscale, the calibrated LUT is reliable. Finally, we find that under the calibrated LUT, phase production range above 2π and the gray-phase linear fit reached 99.83% and 99.72% respectively for 920 nm and 1064nm. The linear phase curve proves that using the calibrated LUT, the SLM can accurately generate the phase mask designed by the GSSIP algorithm without additional errors.

### Performance measures in patterned illumination pattern

We introduce the following quality measures: root-mean-square (RMSE)^7^, modulation efficiency^8^, and uniformity^9^, to evaluate the performance of the GSSIP algorithm for holographic beam shaping. The RMSE is to measure the accuracy of the illumination pattern:

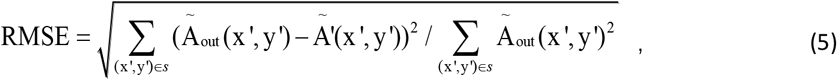

where Ã_*out*_ and Ã′ respectively denote the target intensity distribution and normalized reconstructed illumination pattern. The coordinate (x′, y′) is the position on the output plane. Symbol *s* represents the pixels in the signal area where the illumination pattern is addressing. The modulation efficiency represents the light energy utilization of the modulation by the phase mask, which is the ratio of the power in the signal region to the total power in the output plane:

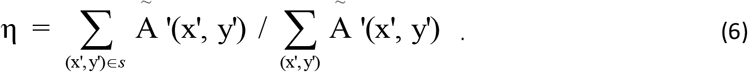

For two-photon holographic fluorescence imaging, the modulation efficiency could not be directly obtained from fluorescence images, but the SNR of the images is positively correlated with the efficiency. The SNR is the ratio of the average intensity 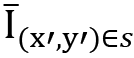 of the signal area to the average intensity 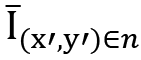 of the noise area:

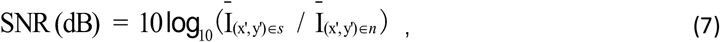

where the noise area is located within 50 μm from the signal area, and the two regions shared the same shape. The uniformity indicates the flatness of the holographic beams and is calculated by:

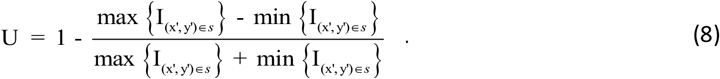

The RMSE, modulation efficiency or SNR, and uniformity are synthetically evaluated the modulation quality of holographic beams, which could more comprehensively characterize the effectiveness of the TF-GSSIP, TF-GS, and other methods for beam shaping.

### Target patterned illumination and data analysis in photostimulation

The illumination pattern used for photostimulation is customized through the GSSIP algorithm. Because photoactivation is conspicuous when the power of excitation light for calcium imaging is relatively weak, the SNR of neurons fluorescence image is low, which is not conducive to the identification of target cells. To overcome the problem, the target holographic pattern of the target cell comes from the binary result of the projection of multiple images. The binary image acts as the target light distribution on the output surface in the GSSIP algorithm. A phase mask is designed by the algorithm and loaded on the SLM. In the photostimulation system, the distribution of femtosecond beams exactly covers the target cell through the phase modulation (Extended Data Fig. 5b).

An image stack of ∼1800 consecutive frames (from GCaMP6s ‘green’ channel) is obtained by the sCMOS during photostimulation. Data analysis is performed using custom-written software in ImageJ (Fiji) and MATLAB. Cellular regions of interest (ROIs) are manually drawn using the maximum intensity projection of the recorded image stack, and mean fluorescence time-courses (GCaMP6s) in each image are extracted. Calcium signals (ΔF/F) of each ROI are calculated according to temporal averaging^10^ and corrected background ^11^. We classified neurons as responsive if they shows a significant difference in the average fluorescence in the 500 ms time window before and after the photostimulation (****p<0.0001, two-sample Student’s t test).

### Hippocampus neurons cultures

P0 rat pups are used for the in vitro hippocampus neuron culture. The 28.2-mm culture dish (Nest, 801001) with 15-mm glass bottom is pre-coated with poly-D-lysine (10ug/ml, Sigma, 32160801) for 20-24 hours and wash three times with water to remove redundant poly-D-lysine. Then, the culture dish is coated with laminin mouse protein (20μg/mL, Gibco, 23017015) for another 24 hours and wash three times with water to remove redundant laminin mouse protein. The residual water is removed by air dry for 30 minutes. About 30000 neurons are seeded on the culture dish on DIV0 and cultur at 37°C with 5% CO2. Culture medium (Neural basal plus, Gibco, A3653401) is supplemented with penicillin (100U/ml), streptomycin (100 g/ml), B27 plus (1:50 dilution, Gibco, A3582801), and GlutaMAX (1:100 dilution, Gibco, 35050087). Neurons are co-infected with C1V1-mCherry AAV and GCaMP AAV on DIV6-8 and imaged on DIV8-20.

### Scattering specimen preparation

Sagittal brain slices are prepared from 6-8weeks old mice. Mice are rapidly anesthetized with pentobarbital sodium [1%, wt/vol, 0.1 mL/10 g, intraperitoneal (i.p.)] and transcardially perfused with ice-cold NMDG solution. The brain is also dissected out under ice-cold NMDG solution [contains (in mM) 92 NMDG, 25 glucose, 5 sodium ascorbate, 2 thiourea, 3 sodium pyruvate, 10 MgSO4, 0.5 CaCl2, 2.5 KCL, 1.25 NaH2PO4, 30 NaHCO3, and 20 HEPES, all chemicals from Sigma] ^12^. Sections of 100 µm, 200 µm, 300 µm, 400 µm, and 500 µm are cut with a vibratome (VT1000S, Leica), using speed 0.16–0.20 mm/s and amplitude of 1.6 mm. Transferred brain slices to HEPES holding solution [contains (in mM) 92 NaCl, 25 glucose, 5 sodium ascorbate, 2 thiourea, 3 sodium pyruvate, 2 MgSO_4_, 2 CaCl_2_, 2.5 KCL, 1.25 NaH_2_PO_4_, 30 NaHCO_3_, and 20 HEPES, all chemicals Sigma] and bubbling with 5% CO_2_-95% O_2_ at 20-24 °C for 1 hour ^12, 13^. We do not use PFA fixation and sucrose dehydration to ensure that the scattering effect of the brain slice is similar to in-vivo. The sections are then affixed to the fluorescent plate (Thorlabs) and used spacers of different thicknesses to prevent the slices from being deformed before mounting the slices. All measurements are completed within 6h after sectioning.

For skull preparation, mice aged 6-8 weeks are anesthetized with sodium pentobarbital and perfused transcardiacally with ice-cold 0.01 M PBS (Solarbio). After decapitation of the perfused mouse, the skull is completely taken out. We use a spiral micrometer to measure the average thickness of the mouse skull to be about 150 µm ^14, 15^. The skull is ground into three circular scattering media with a diameter of 3 mm and an average thickness of 150 µm, 100 µm, and 50 µm, respectively. To ensure that the skull scattering effect is similar to in-vivo, various tests should be completed within 6h.

---

**Extended Data Fig 1.**
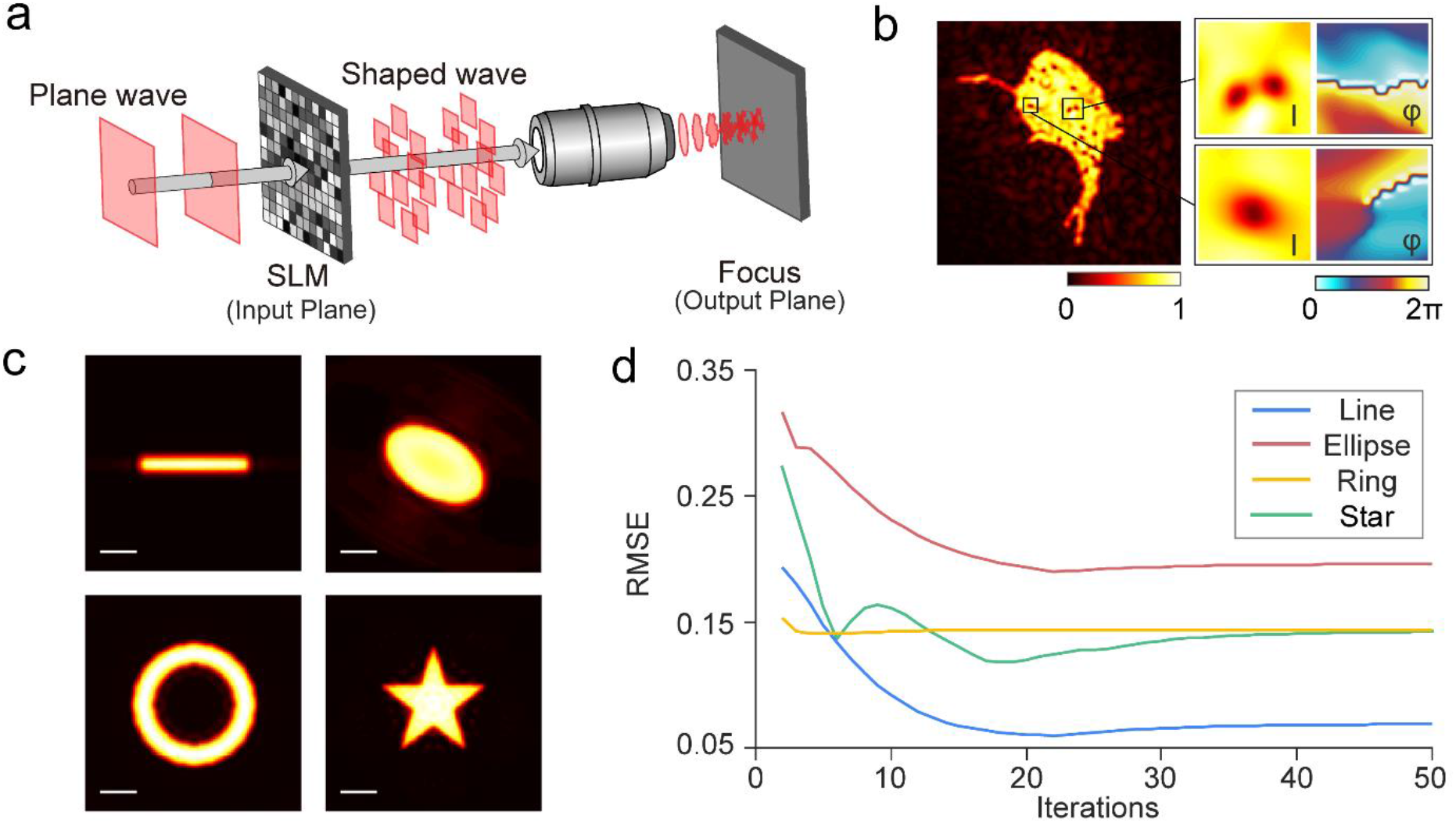
Optical configure for CGH and some features of the GSSIP algorithm. (**a**) The schematic diagram for the CGH. The input and output planes are located at the front and back focal planes of an objective. The plane wave is modulated to the target-shape illumination in the holographic domain. (**b**) Some examples of intensity fluctuations in patterned illumination designed by the GS algorithm. Each collection contains amplitude (upper row) and phase (lower row) in the form of intensity images (left column) and 3D plots (right column). (**c**) Representative patterned illumination designed by the GSSIP algorithm in numerical simulation: a 1-D line, a non-angular ellipse, a hole-shaped ring, and a concave-polygon star. (**d**) The curve of RMSE varying with the number of iterations in the GSSIP algorithm for designing the target-shaped illumination (shown in c), showing that GSSIP algorithm converges after about 20 iterations. Scale bar 15 μm.

**Extended Data Fig 2.**
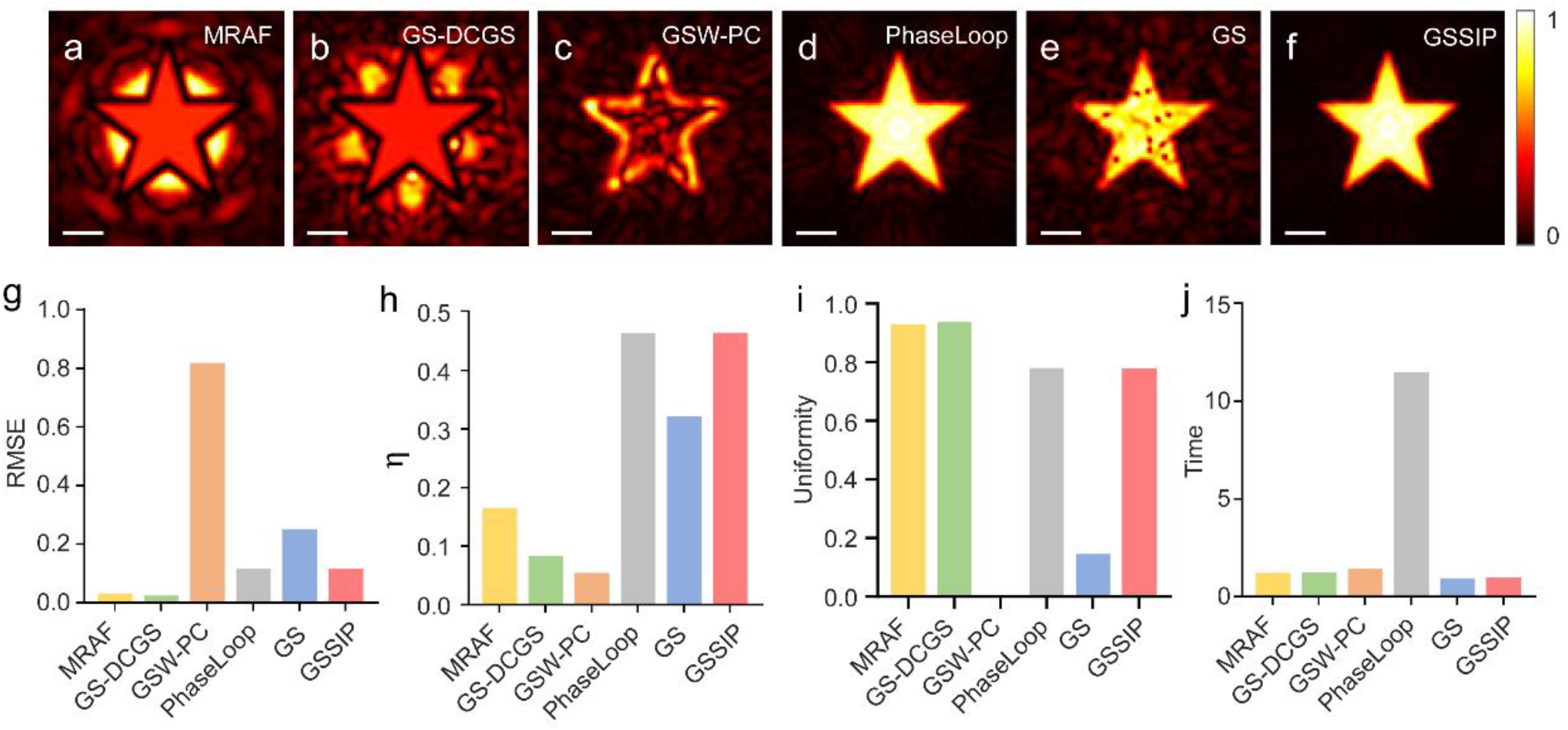
Comparison between the GSSIP algorithm and other speckle-free CGH algorithm based on designing 2-D star-shaped illumination. (**a-f**) Predicted patterned illumination output by MRAF, GS-DCGS, GSW-PC, Phase- Loop, GS and GSSIP algorithm in the numerical simulation. (**g-j**) The RMSE, modulation efficiency (η), uniformity and algorithm-running time of illumination patterns designed by different algorithm (shown in Figure. a-f). Scale bar 15 μm.

**Extended Data Figure 3.**
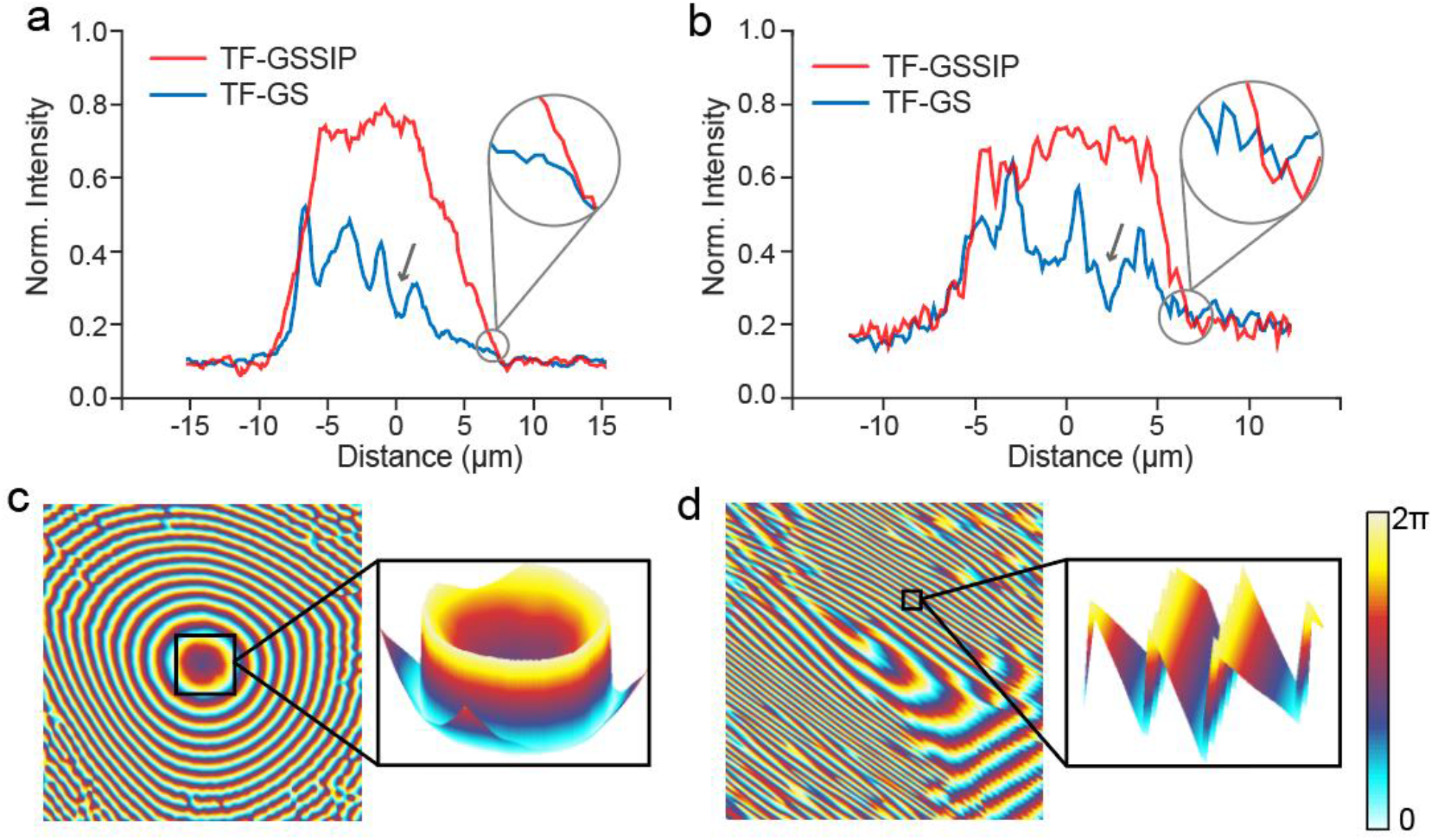
Intensity distribution and phase mask in TF-GSSIP for a neuronal soma and a dendrite. (**a**,**b**) Intensity profile of the neuronal soma and the dendrite shown in Figure 1d,e,g,h. White arrows indicate the direction of profiles. It is obvious that the intensity distribution (red trace) is with sharp edges (zoomed insets) and the intensity is enhanced and uniform. Both features indicates the profile is superior to the results of TF-GS (blue trace), showing a slow- reduction edges and notable intensity fluctuations. (**c, d**) Phase masks designed by the GSSIP algorithm, corresponding to the neuronal soma and the dendrite shown in Figure 1e, h. The magnification boxes present 3-D distribution of phase masks and verify the spatially continuous wavefront modulation.

**Extended Data Fig 4.**
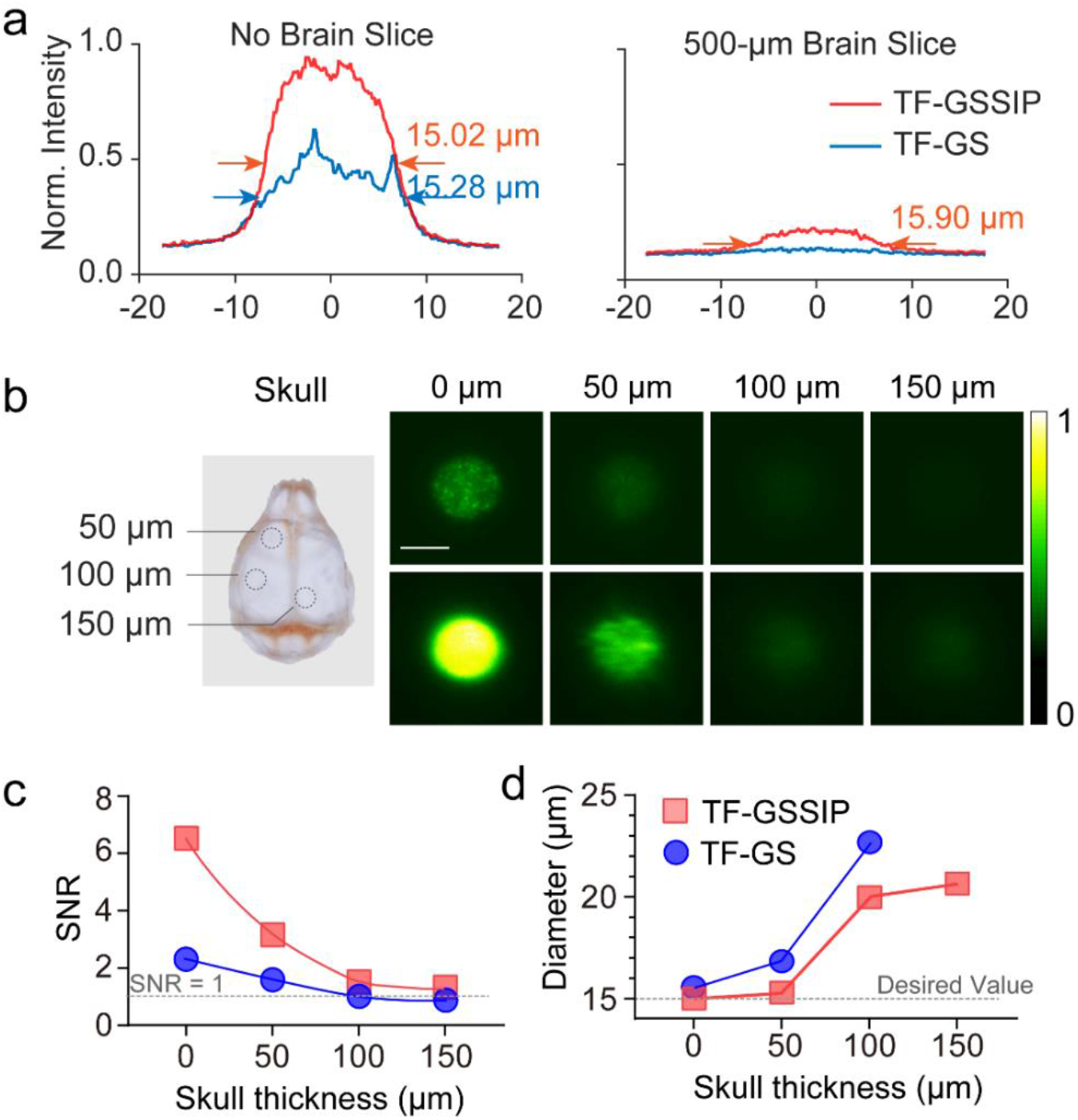
Intensity distribution after penetrating brain slices and patterned focusing after penetrating mouse skull. (**a**) Lateral profile of the 15-μm-diameter spot scattered by none or a 500-μm-thickness brain slice (shown in Figure. 2b). It is clear that the FHWM of TF-GS (blue curve) is larger than that of TF-GSSIP (red curve) (left panel). A visible envelope of the intensity profile (red curve in the right panel) can be observed when the TF-GSSIP patterned illumination focused through a 500-mm-thick slice with diameter (defined as the FHWM of the intensity profile) varying less than 1 μm, while there is not a recognized intensity profile in TF-GS (blue curve). (**b**) 2P fluorescence images of a 15-μm-diameter holographic spot generated by TF-GS (top) and TF-GSSIP (bottom) after passing through fixed skulls (0, 50, 100, and 150 μm) at the laser power of 0.10 mW/μm2. (**c, d**) SNR (Figure c) and Uniformity (Figure d) of the conventional and TF-GSSIP patterned foci shown in Extended Data Fig. 4b as a function of skull thickness. The diameter of holographic spot by TF-GS after passing through the 150-μm-thickness skull is not shown, because the fluorescence signal is not visible (shown in Extended Data Fig. 4b, and SNR ∼ 1 shown in Extended Data Fig. 4c).

**Extended Data Fig 5.**
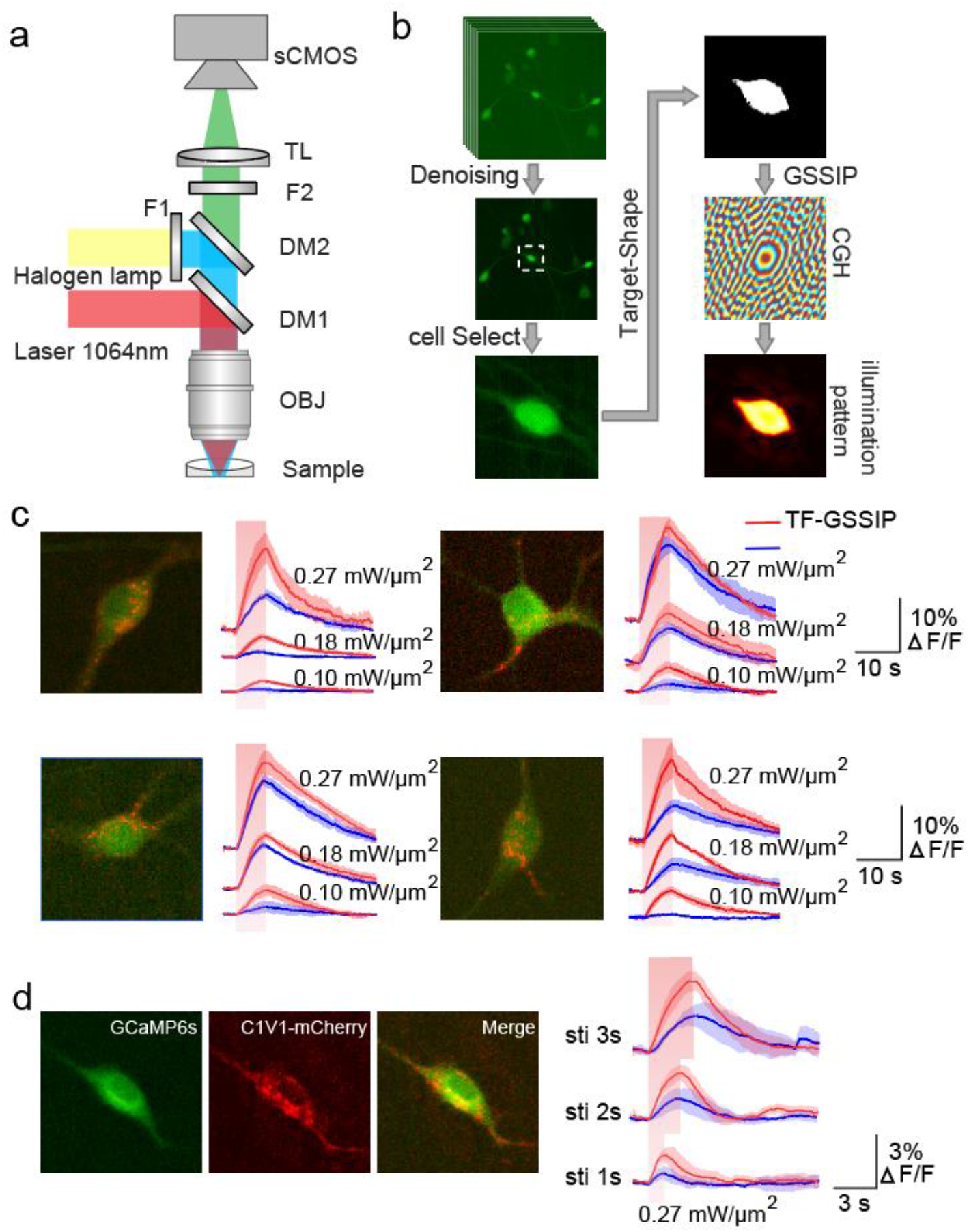
Experimental implementation for photostimulation and calcium responses to different stimulus power and duration. (**a**) Schematic diagrams for our all-optical recordings and perturbation system. F, Filter;DM, dichroic mirror; OBJ, objective; TL, tube lens. (**b**) Flow chart of the experiment procedure, which includes five steps: image denoising, target-neuron identification, cell body extraction, phase masks calculation, and the final patterned illumination genetated by TF-GSSIP. (**c**) Calcium responses of 4 cells stimulated by TF-GSSIP (red trace) and TF-GS (blue trace) system upon different stimulation power, which are counted in Figure. 5f. (**d**) An example of calcium responses recordings during 1, 2, and 3 s photostimulation in TF-GS and TF-GSSIP system.

**Extended Data Table 1.**
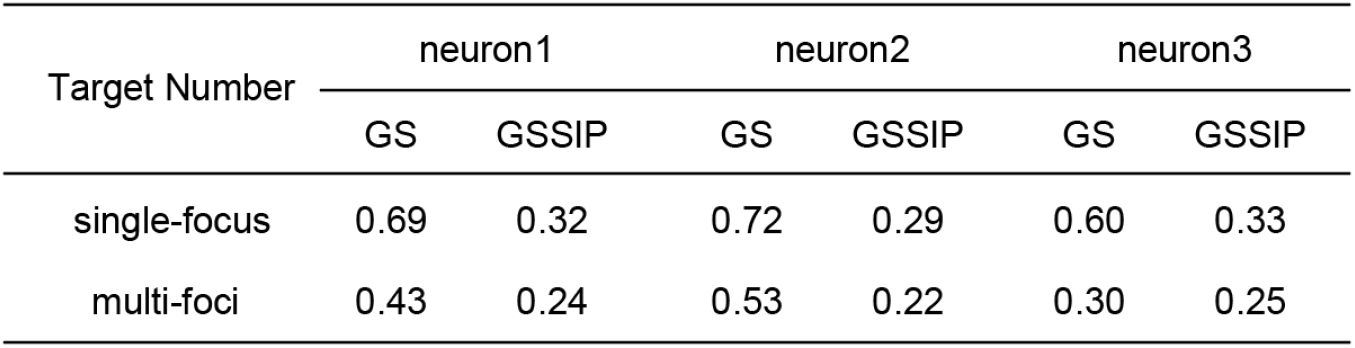
RMSE in the single-target and multi-target patterned illumination shown in Figure. 3c and Figure. 3e.

| Target Number | neuron1 |  | neuron2 |  | neuron3 |  |
| --- | --- | --- | --- | --- | --- | --- |
|  | GS | GSSIP | GS | GSSIP | GS | GSSIP |
| single-focus | 0.69 | 0.32 | 0.72 | 0.29 | 0.60 | 0.33 |
| multi-foci | 0.43 | 0.24 | 0.53 | 0.22 | 0.30 | 0.25 |

